# A simple geometrical model of the electrostatic environment around the catalytic center of the ribosome

**DOI:** 10.1101/2022.03.07.483325

**Authors:** Marc Joiret, Francesca Rapino, Pierre Close, Liesbet Geris

## Abstract

The central function of the large subunit of the ribosome is to catalyze peptide bond formation. This biochemical reaction is conducted at the peptidyl transferase center (PTC). Experimental evidence shows that the catalytic activity is affected by the electrostatic environment around the peptidyl transferase center. The electrostatic field originates from the particular 3-dimensional space distribution in the phosphate moieties of the 23S (archea and bacteria) or 28S (eukarya) rRNA of the large ribosomal subunit as well as the funnel shape of the cavity at its connection with the ribosome exit tunnel. Here, we set up a minimal geometrical model fitting the available x-ray solved structure of the ribonucleoproteic cavity around the catalytic center of the large subunit of the ribosome. The purpose of this phenomenological model is to estimate quantitatively the electrostatic potential and electric field that are experienced during the peptidyl transfer reaction. At least two reasons motivate the need for developing this quantification. First, we inquire whether the electric field in this particular catalytic environment, made only of nucleic acids, is of the same order of magnitude as the one prevailing in catalytic centers of the proteic enzymes counterparts. Second, the protein synthesis rate is dependent on the nature of the amino acid sequentially incorporated in the nascent chain. The activation energy of the catalytic reaction and its detailed kinetics are expected to be dependent on the mechanical work exerted on the amino acids by the electric field, especially when one of the four charged amino acid residues (R, K, E, D) is newly incorporated in the nascent chain. Physical values of the electric field will provide quantitative knowledge of mechanical work, activation energy and kinetics of the peptide bond formation catalysed by the ribosome.

## 1. INTRODUCTION

Ribosomes are the cells’ manufacturing tools for building up proteins. They decode the 61 sense codons from a primary message encrypted in a messenger RNA (mRNA) single molecule. They translate it with the help of a set of transfer RNAs (tRNAs) into 20 amino acids to be sequentially polymerized in a nascent polypeptide that will eventually pass through the ribosomal exit tunnel and fold into its final structure.

X-ray solved structural representations of ribosomes have been publicly available for different species at atomic resolution for more than 20 years. The peptide bond is formed between the nascent protein chain and a newly incorporated amino acid at the ribosomal large subunit catalytic center, whose salient feature is that the peptidyl transferase center (PTC) is not an enzyme but a ribozyme, composed of ribosomal RNA (rRNA). There are no ribosomal protein components within a 15 *Å* radius of the catalytic center [1, 2]. The electrostatic environment of the PTC is largely determined by the presence of the phosphate moieties belonging to the 23S rRNA backbone in archae or bacteria, and to the 28S rRNA backbone in eukarya.

Efforts in understanding the kinetics of the protein synthesis in vivo and the factors that affect elongation rate have been conducted for decades by the research community [3, 4]. Experimental evidence supports the fact that enzymatic kinetics is affected by the electrostatic environment around the active sites of enzymes. Recently, vibrational Stark effect spectroscopy techniques were used to obtain direct measurements of electric fields at active sites of enzymes [5]. To elucidate the mechanisms of action and the reaction kinetics, there is great interest in knowing quantitatively the potential and the electric field prevailing at the active site of the ribosome, at the peptidyl transferase center and along the ribosome exit tunnel. To this date, there is no experimental protocol that would bring the large subunit of the ribosome amenable to such direct measurements, although pioneering efforts were undertaken in the ribosome exit tunnel of rabbit reticulocytes [6].

Given the lack of direct experimental measurements for the potential and the electric field, the only way to obtain the physical quantities of interest is through the use of a mathematical model. Such mathematical models are broadly used to shed light on structure, function and properties of often complex biomolecular systems and are used in numerous studies [7]. In this paper, we start from the publicly available atomic structure of the ribosome and use the fundamental laws of electrostatics along with the dielectric properties of media inside the ribosome to calculate the potential and the electric field experienced around the center of the peptidyl transferase reaction.

## 2. MATERIAL AND METHODS

### 2.1. X-ray crystallographic space position of phosphate moieties and charged amino acid residues around the PTC cavity

We analyzed the publicly available structure of the large ribosomal subunit of the archeon *Haloarcula marismortui* (PDB code: 4V9F downloaded from https://www.rcsb.org/) obtained from x-ray crystallography at 2.4 *Å* [8]. To find the ribosome tunnel entry port and extract the atom coordinates around the PTC, we used a tunnel search algorithm developed by Sehnal *et al* [9], implemented in MOLE 2.0 and the web-based MOLEonline 2.0 tool publicly available online [10, 11]. We used PyMOL (PyMOL Molecular Graphics System, Version 2.3.2) and exported the relevant selected atom positions’ cartesian coordinates to output files. These files were further processed with algorithms coded in Python to select the charged chemical groups on or near the inner surface of the PTC cavity. To localize the PTC, we used the geometric center of the five known 23S rRNA P loop and A loop nucleotides that are known to interact with the 3’-terminal CCA end of the tRNA acylated to the carboxy terminal amino acid of the nascent chain at the P site or the 3’-terminal CCA end of the aminoacylated tRNA at the A site [2, 12]. These five nucleotides in the P and A loops of 23S rRNA are A2485(2450), A2486(2451) for the P loop and G2588(2553), U2589(2554) and U2590(2595) for the A loop [1]. The numbering of the 23S rRNA residues is based on the *H. marismortui* sequence, while the corresponding position in the *E. coli* ribosome is shown in the brackets. We translated the crystallographic data model space so that the tunnel entry point would be at the origin and we aligned the direction from the tunnel entry point to the PTC geometric mid-point along the positive z-axis. We selected all charged amino acid residues (NH2 or NZ for arginine or lysine, OE2 or OD2 for aspartate or glutamate) belonging to ribosomal proteins and all charged non-bridging oxygen atoms bound to the phosphate moieties in the 23S rRNA backbone that are closer than 40 *Å* from the centerline joining the tunnel entry port to the PTC geometrical midpoint. The cavity around the PTC was approximated by fitting a truncated prolate spheroid (ellipsoid of revolution about the major axis) having its semi-major axis aligned with the direction from the tunnel entry port to the PTC. The half prolate spheroid was also scaled in such a way that its semi-major axis spans the distance between the tunnel entry point and the decoding center P site (∼ 8.75 nm) and its semi-minor axis spans a half-length of 3 codons (∼ 4.50*/*2 = 2.25 nm). We algorithmically set out the 3D equations of the truncated prolate spheroid in this reference frame to calculate the radial distance of the selected atoms to the surface of the prolate spheroid PTC cavity. We confirmed that there are no charged atoms belonging to the 23S rRNA or to any ribosomal proteins inside the fitted truncated prolate spheroid cavity.

## 3. THEORY AND CALCULATIONS

### 3.1. Idealized shape model of the ribosomal RNA cavity around the PTC and the Yukawa-Debye-Hückel potential with dielectric screening

After the sphere, the prolate spheroid is the geometrical shape with the smallest surface encompassing the largest volume. An electrostatic model of the cavity around the catalytic center of the large subunit of the ribosome is built using this most simple shape. This fulfills the minimal geometrical constraints which prevail between the ribosome peptide exit tunnel, the mRNA channel and the size of three aminoacylated-tRNAs that are accommodated in the cavity during nascent protein elongation Fig.1(a). The prolate spheroid was fitted onto the aforementioned publicly available x-ray solved structure of the large ribosomal cavity around the peptidyl transferase center as explained in Material and Methods.

**FIG. 1:**
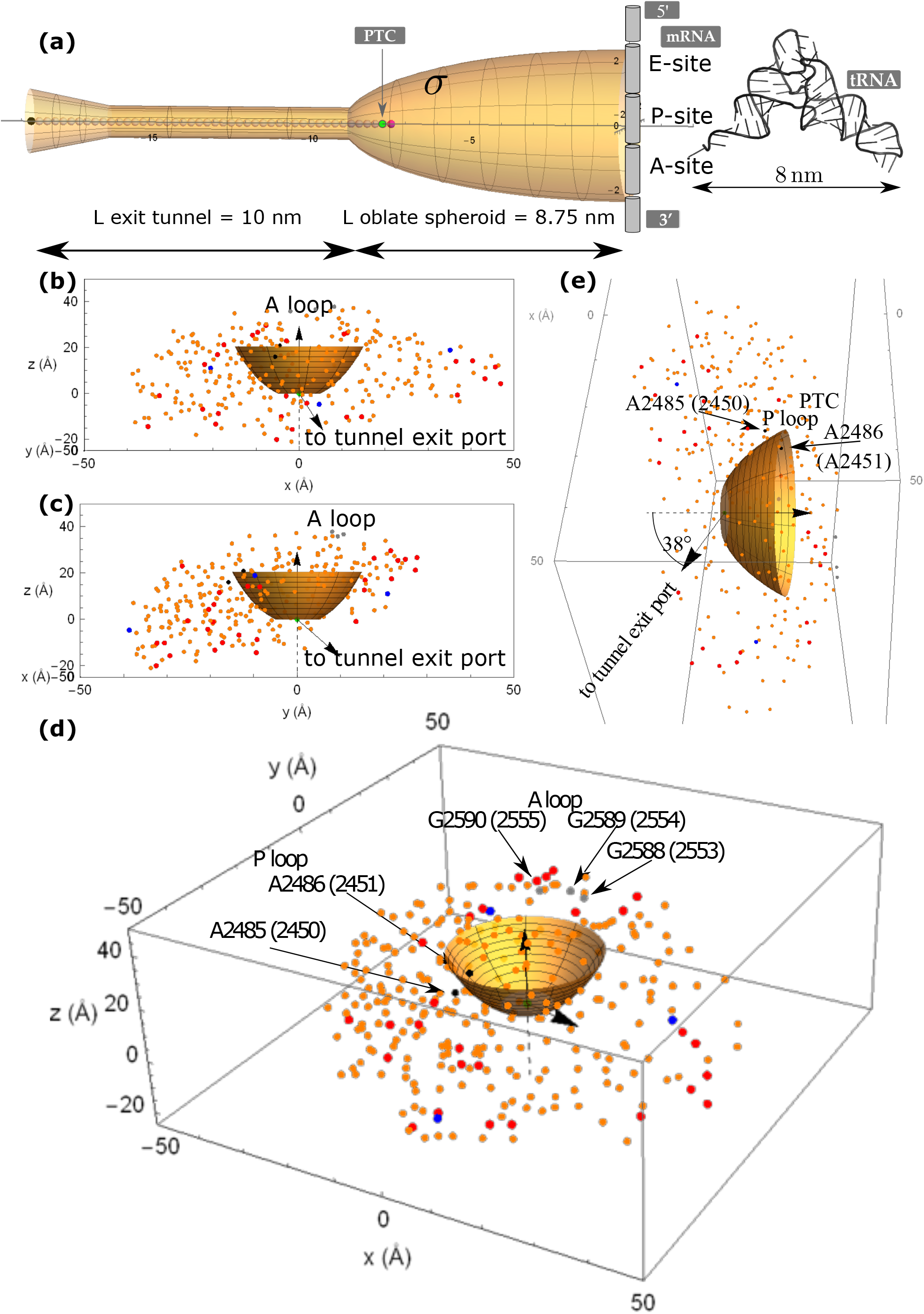
(caption of next page).(a) Ribosome exit tunnel and cavity around the PTC. (b) and (c) Front and right view of the charged groups neighbouring the PTC cavity. (d) and (e) 3D scatter plot of the 291 charged atoms around the PTC. Phosphorus atoms bridging the 23S rRNA backbone (orange dots), positively charged moieties of lysine or arginine belonging to ribosomal proteins (red dots), negatively charged moieties of aspartate or glutamate belonging to ribosomal proteins (blue dots), phosphorus atoms of P loop 23S rRNA nucleotides A2485 and A2486 (black dots), phosphorus atoms of A loop 23S rRNA nucleotides G2588, U2589 and U2590 (gray dots). The atomic positions were retrieved from *H. marismortui* x-ray solved structure of the large subunit of the ribosome.

The electrical scalar potential 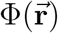 at the observed position 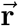 is expressed, in a homogeneous medium of constant permittivity and in the absence of dielectric screening effects, by the Coulomb law:

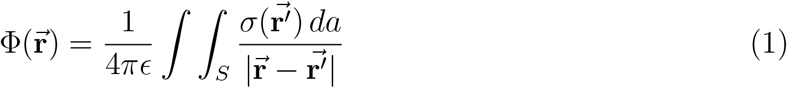

where 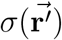 is the surface-charge density (measured in coulombs per square meter) at position 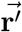 of the source, *da* is the two dimensional surface element at 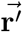 and *∈* is the permittivity of the dielectric medium (Eq. 1.23 in Jackson [13]) with *∈* = *∈*_*r*_*∈*_0_, where *∈*_*r*_ is the relative permittivity of the medium and *∈*_0_ is the permittivity of free space.

It is worth emphasizing here that the position where the physical values of the potential is calculated is 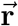, but the variable over which the surface integration is conducted is 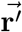 with its elementary surface elements 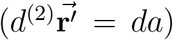 integrated over the surface domain of interest in the 3D space. We actually sum over all the fixed charges located on the surface. There are two ways to describe the charges and their positions. In the first, the charges are considered to be continuously distributed on the surface. A surface charge density *σ* must be known which can be a function of 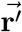 as expressed in equation (1). In the second, the real fixed charges are discrete (with no spatial extensions) but the surface domain on which integration is conducted is still a compact interval in ℝ^2^. Implicitly, discrete charges *q*_*i*_, spatially localized in 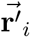 are mathematically represented by a generalized function which is the product of the charges *q*_*i*_ by a Dirac distribution [14]: 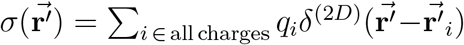. This second type of description will be used in the next section. In mathematical terms, in both descriptions, the integration domain is the support of the function to be integrated. The support of the function is the interval upon which the function exists and is not null. 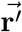 lives in the support of the charges or the support of the Dirac functions that you implicitly use to define the position of the charges that are sources of the field. 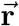 lives in another space, the set of points where you actually want to calculate the field, irrespective of the positions of the source charges. In the surface integral calculation conducted below, 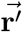 has coordinates *u, v* in the support of the charges, which is a surface in 3D (a half prolate spheroid) and 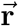 has the cartesian coordinates (0, 0, z) of a straight line in 3D, because for the sake of simplicity, the potential and the electric field will only be calculated along the central axis *z*.

In the presence of polarizable dielectric material, screening effects occur due to induced dipoles in the medium separating the fixed charged sources of the field and the observation point. In this latter case, the electrical scalar potential 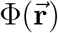 at the observed position 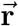 is expressed by the Yukawa-Debye-Hückel law:

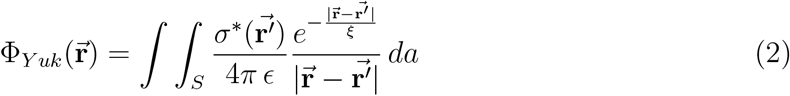

where 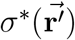 is the actual *formal bare* surface charge density. There is a marked exponential damping of the Coulomb interaction where *ξ* is a characteristic distance of the screening. We assume the screening effect to be homogeneous around the fixed charged surface, and thus we take the assumption that both *∈* and *ξ* do not depend on 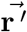 or 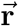 or at least are piecewise constants in a given space domain [15].

For the prolate spheroid, we can take advantage of the axial symmetry and restrict the observation positions to the spatial points on the *z* axis, i.e., for 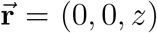. The surface integration is conducted on the support of the fixed source charges. The prolate spheroid’s inner wall is geometrically generated by the *γ*(*u*) curve moving axially along the z-axis from *z* = 0 to *z* = −*a* as drawn in Fig.1(a) and inset in Fig.2(a), where *a* and *b* are the semi-major and semi-minor axis of the prolate spheroid:

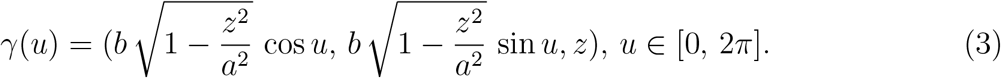

**FIG. 2:**
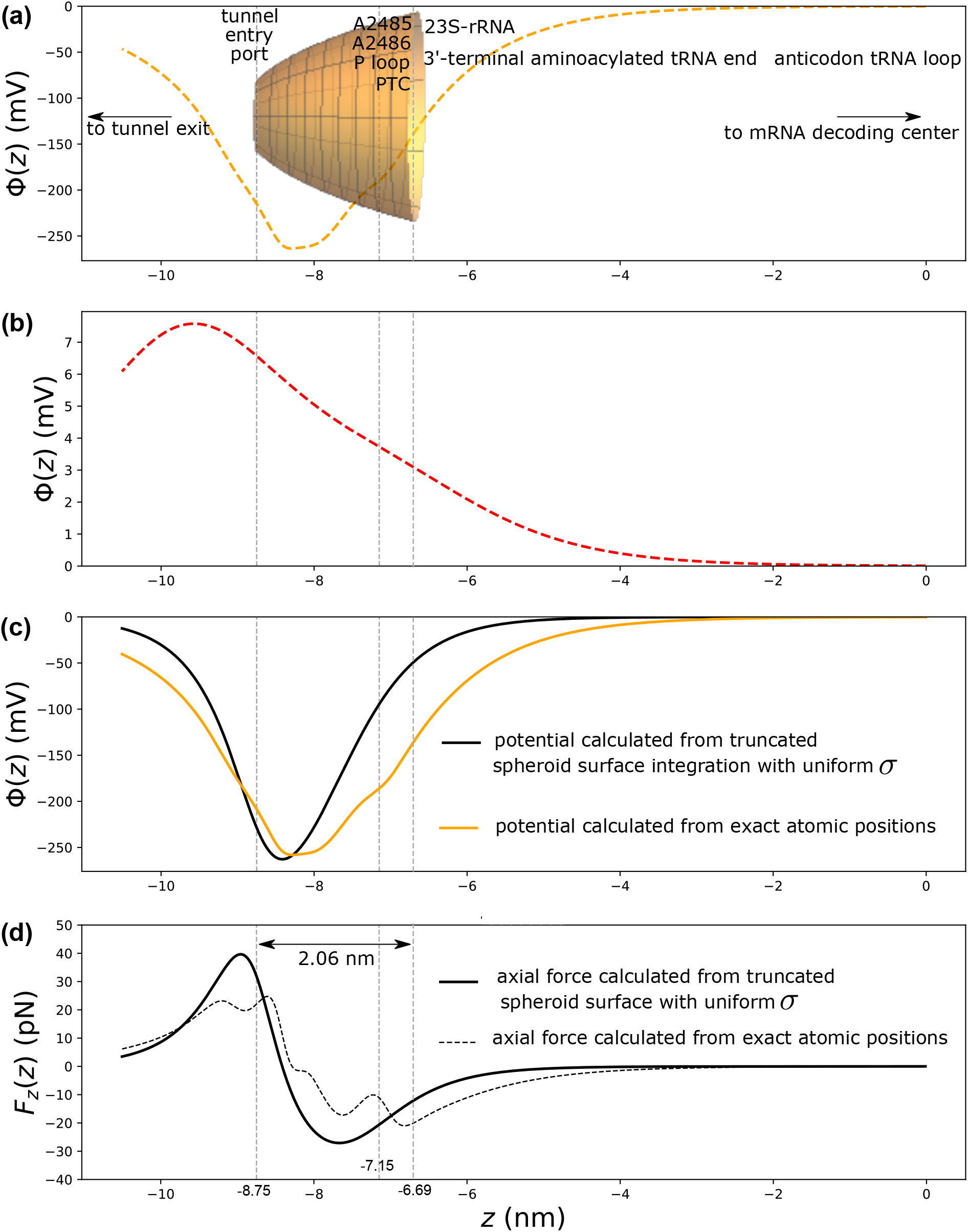
Electrostatic potential profiles contributed by the ribosomal large subunit cavity around the PTC. (a) Electrostatic potential (mV) computed from the exact atomic positions of the 100 phosphorus within 12 *Å* of the surface cavity. (b) Electrostatic potential (mV) computed from the exact atomic positions of the 13 charged amino acid within 12 *Å* of the surface cavity. (c) Electrostatic potential (mV) computed from the exact atomic positions of all 113 charged moieties within 12 *Å* of the surface cavity (orange solid) compared to the potential resulting from a uniformly charged truncated spheroid surface of *σ*\* = −6.35|*e*|*/*nm^2^ (black solid). (d) Axial electrostatic force (pN) computed from the exact atomic positions of all 113 charged moieties within 12 *Å* of the surface cavity (gray dashed) compared to the force resulting from a uniformly charged truncated spheroid surface of *σ*\* = −6.35|*e*|*/*nm^2^ (black solid). All panels: gray dashed vertical lines, from left to right: tunnel entry port position (radius= 5 *Å*), P loop A2485 z position, truncated spheroid position where the cavity radius = 15 *Å*. A Bjerrum screening length of 0.72 nm was used for the analytical potential calculation and a Debye screening length of 1.1 nm was used in the discrete sum over the exact atomic positions, with ionic strength *I* = 75 mM*/*L. The water permittivity *∈* = 78 was assumed inside the PTC cavity.

The prolate spheroid’s half surface can be written as *S* = *ϕ*(*K*) where 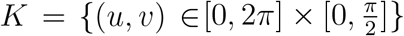 and where *ϕ* : ℝ^2^ → ℝ^3^ :

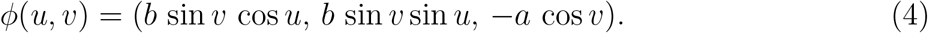

Another equivalent parametrization of the prolate spheroid’s half surface can be written as *S* = *ϕ*(*K*) where *K* = {(*u, v*) ∈ [0, 2*π*] × [0, 1]} and where *ϕ* : ℝ^2^ → ℝ^3^:

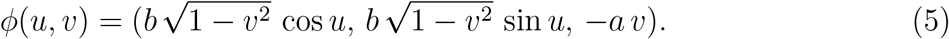

Upon these parametrizations, the distance between any observation point and any point supporting a source charge can be expressed respectively as:

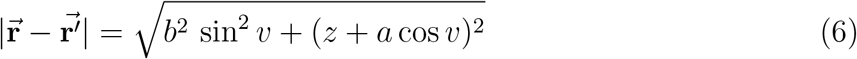

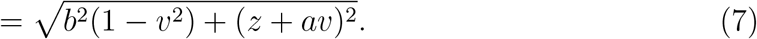

There is an aperture at the leftmost end of the prolate spheroid where the PTC cavity is connected to the ribosome exit tunnel which is approximated by a cylinder as shown in Fig.1(a). The radius of the aperture is around 5 *Å*. Hence, the exact support of the resulting truncated spheroid’ surface is parametrized as *S* = *ϕ*(*K*) where *K* = {(*u, v*) ∈ [0, 2*π*] × [*v*_*inf*_, *v*_*sup*_]} where *v*_*sup*_ *<* 1. *D*_*u*_*ϕ* is the first partial derivative of the parametric equation of the surface *ϕ*(*u, v*) with respect to *u* and *D*_*v*_*ϕ* is the first partial derivative of the parametric equation of the surface *ϕ*(*u, v*) with respect to *v*. In the general Eqs (1 and 2), the surface-charge densities 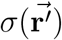 or 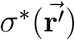 are dependent of the position 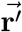 on the support of the source charges. Here, we will take the simple approximation that *σ* or *σ*\* can be considered a constant parameter over a surface of a given shape, e.g. over the spheroid’s surface. This is the surface charge uniform distribution assumption for a given shape. Note however that a space dependence of *σ* is possible if it is compensated for by a similar space dependence of *∈* ensuring the combined ratio *σ/∈* is constant in a region of interest. This piecewise constant ratio is the strictly necessary assumption for the mathematical surface integration calculations of our models to be analytically tractable. In subsection 4 4.2, we will discuss how reasonable this assumption is by comparing the potential profiles calculated from the exact x-ray crystallographic structural data of the large subunit of the ribosome and calculated from the idealized spheroidal shape with constant surface charge density and a constant medium permittivity.

The electrostatic scalar Yukawa screened potential (2) results from the surface integral calculation:

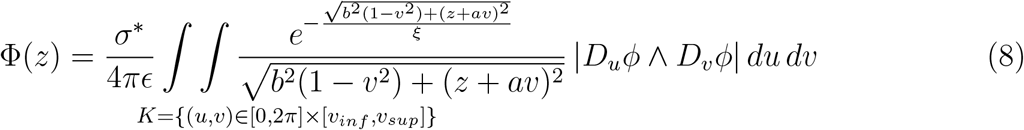

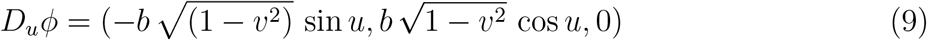

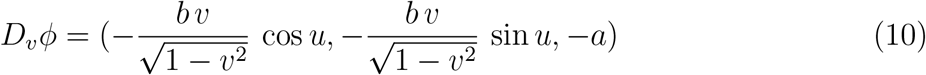

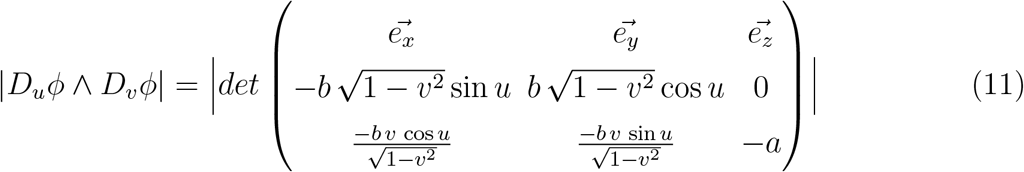

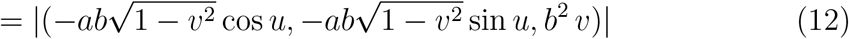

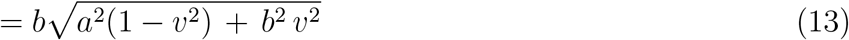

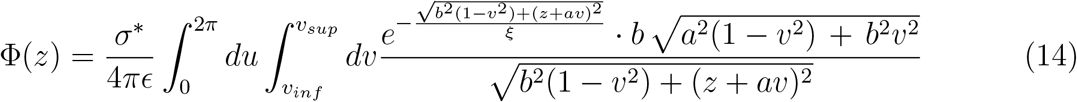

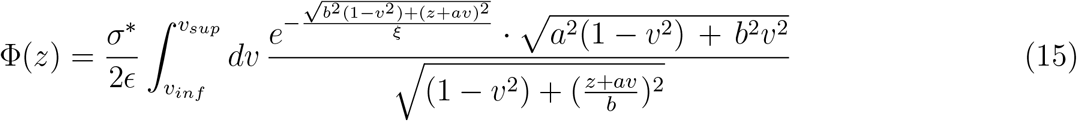

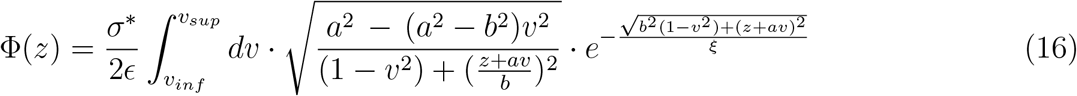

The integral converges if the denominator in the integrand of Eq.(16) does not vanish, i.e., if *v*_*sup*_ *<* 1, meaning that the spheroid (ellipsoid of revolution) is truncated. The ribosome cavity around the PTC further goes towards the ribosome exit tunnel which is approximated by a cylinder surface. By virtue of continuity and physical consistency, it is necessary that the electrostatic potential be a continuous function of space inside the cavity of the catalytic center of the ribosome and inside the ribosome exit tunnel.

According to the x-ray solved structure of the ribosomal large subunit, the length scales for the spheroid are with semi-major axis *a* = 8.97 nm, semi-minor axis *b* = 2.25 nm, giving *v*_*sup*_ = 0.975 such that the aperture of the spheroid cavity towards the cylindrical ribosome exit tunnel has a radius *R* = 5 *Å*. The length between the center of the spheroid, at *z* = 0, and the entry point of the ribosome exit tunnel is 8.75 nm. The P loop 23S rRNA nucleotide A2486 (2451) interacting at the P site in the peptide bond formation has its axial position 20.6 *Å* away from the tunnel entry port. This provides the lower integration limit for the support of the charges on the spheroid’s surface *v*_*inf*_ = 0.74536.

The integral in Eq.(16) converges but cannot be solved analytically as it is an elliptical integral. We resorted to two classical numerical approaches using Newton-Cotes method or Gaussian quadrature where the nodes sampling followed the Gauss-Konrod rule[16]. Both methods gave the same result for the solution of the electrostatic potential Φ(*z*) profile along the centerline of the truncated spheroid’s cavity.

The axial electric field is obtained from the negative of the first partial derivative with respect to *z*:

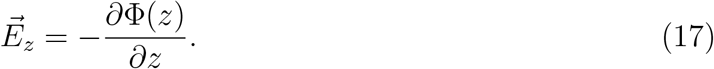

Finally, the axial force on a test charge can be calculated from:

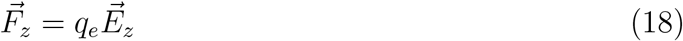

where *q*_*e*_ is the charge of the test charge.

In Eq.(16), we require the knowledge of at least two phenomenological parameters: the *formal bare* surface charge density *σ*\* on the truncated spheroid surface in the vicinity of the PTC and the dielectric response (permittivity) *∈* of the medium around the PTC. In fact, only the ratio *σ/∈* is needed. The phenomenological screening length parameter *ξ* is also required.

### 3.2. Structural data model of the ribosomal RNA cavity around the PTC and the Yukawa-Debye-Hückel potential with dielectric screening

The Coulomb or Yukawa-Debye-Hückel electrostatic potential can be calculated from the x-ray solved exact distribution of the source charges (phosphate moieties and charged amino acids) for which the positional 3D map is shown in Fig.1(b) to (e). The method to compute the electrostatic potential based on the real observed atom positions involves a discrete sum and complies with the superposition principle due to the linearity of the electrostatics equations. The Yukawa-Debye-Hückel potential is used and the exact positions 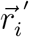 of the sources and their charges *q*_*i*_ are summed over all source charges:

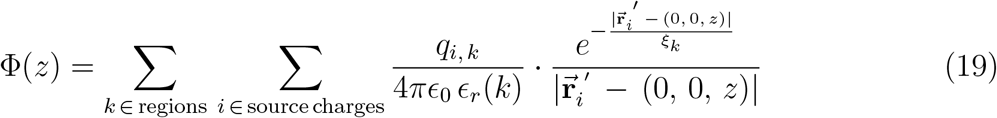

In this formula, two phenomenological parameters are required which are *∈*_*r*_, the relative permittivity of the medium and the screening length *ξ*. The screening length *ξ* is a generic placeholder which can be computed from one of three possible different approaches (Bjerrum, Gouy-chapman or Debye screening length). The Coulomb potential is a particular case of the Yukawa potential when the screening length goes to infinity.

The assumption is made that the two phenomenological parameters are constant (homogeneous) in the media where the potential is computed. The standard or defaulted homogeneous values of these parameters are *∈*_*r*_ = 78 (water) and *ξ* = 10 *Å*. Eq.(19) neglects surface charge polarization effects at dielectric media discontinuities.

In the above Coulomb-Yukawa electrostatic potential Eq.(19), different values of *∈*_*r*_(*k*) and different *ξ*_*k*_ screening lengths can be used in the different *k*-indexed regions (or media). The elementary unit charge value of +|*e*| or −|*e*| = −1.602 10^−19^ C is used for each of the charges *q*_*i*_ associated to the positively or negatively charged atoms at their given 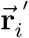 positions.

### 3.3. Phenomenological parameters: *σ*\**/E*, media permittivities and dielectric screening lengths *ξ*

Electrostatic interactions in ribonucleoprotein structures are potentially quite strong, but these interactions are mitigated by the screening effects of water, nucleic acids (both rRNA and tRNAs) or nearby protein atoms [17], even in the absence of mobile ions. In addition to the electrostatic screening, surface charge polarization effects also occur due to dielectric response discontinuities at media interfaces [15].

The screening of electrostatic interactions results primarily from electronic polarization, reorientation of dipolar groups in the vicinity of charges and dipoles. These effects are well understood and can be accurately determined for interactions in isotropic, homogeneous media. However, in complex inhomogeneous environments such as those near the surface of ribonucleoproteins, dielectric screening is difficult to predict. Two factors are expected to be especially important in the case of the ribosome cavity around the PTC and the exit tunnel: the confined geometry and composition of the inner wall close to the surface, and whether the interactions involve direct charges or dipoles. The x-ray solved atomic space positions in the immediate 12 *Å* vicinity of the tunnel wall show that water molecules in addition to tRNAs contribute to the screening of the *formal bare* charges carried by the non-bridging oxygen atoms bound to the phosphorus atoms of 23S/28S rRNA. Due to the dissociation (charge regularization) of surface groups, the rRNA phosphate moieties support surface acquires a net surface charge density that we call *σ*\*. These *bare* charges do not stay unbalanced due to a screening effect involving water solvent. The water molecules dipole moments re-orient so that a layer of positively charged hydrogens oppose the negatively charged phosphate moieties.

Water as a bulk solvent has a relative electric permittivity of 78 (25°C), while experimental and theoretical evidence suggest that proteins (or the nascent polypeptide) have an average dielectric response that can be approximated with a dielectric constant of about 3-4, see [18] and references therein. The dielectric constant of nucleic acids in bulk solution has been measured to be around 8 [19]. Thus, depending on the abundance of water molecules in the PTC cavity volume, the presence of tRNAs and the presence of the carboxy terminal end of the growing nascent peptide, the cavity micro-environment cannot be viewed as uniform. The dielectric constants *∈* should be used in a range from *∈* = 3 − 4 (polypeptide) to *∈* = 78 (water).

Selecting the most appropriate screening theory reduces to knowing which length scale parameter *ξ* to use. Three length scales, i.e., the Bjerrum length (*λ*_*B*_), the Debye length (*λ*_*D*_ = *κ*^−1^) and the Gouy-Chapman length (*ξ*_*GC*_) deserve specific attention as highlighted by Van Roij [20]. The expressions of the Bjerrum, Debye and Gouy-Chapman screening lengths are respectively:

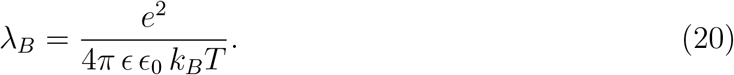

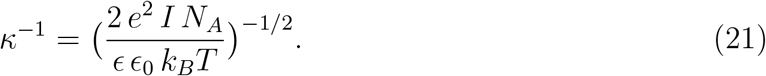

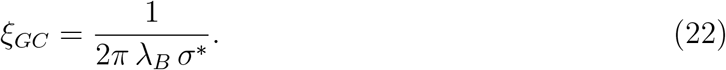

where *e* is the elementary charge of the electron, *∈* the relative permittivity of the medium, *∈*_0_ the permittivity of the vacuum, *I* the ionic strength, *k*_*B*_ the Boltzmann constant, *N*_*A*_ the Avogadro number, *T* the absolute temperature and *σ*\* is the *bare* surface charge density of the wall.

## 4. RESULTS AND DISCUSSIONS

The ribosome exit tunnel was modelled by a cylinder concatenated to a cone frustum in reference [15]. As a result of the scaling exposed in Material and Methods, the peptidyl transferase center (PTC) is approximately surrounded by a minimal surface prolate spheroidal volume accommodating for 3 tRNAs having their anticodon matching the 3 codons at the A, P and E site respectively. The distance between the amino acid residues acylated to their cognate tRNAs and their anticodon loops is 8 nm (the size of the L-shaped tRNA single molecule). The cylinder radius is 5 *Å* [6, 15] and is connected to the PTC cavity by the truncated prolate spheroidal volume as shown in the simplified representation of Fig.1(a). The angle between the support of the PTC arrow and the direction from the tunnel entry to the tunnel exit port is 38°.

### 4.1. Surface charge density and dielectric screening lengths around the PTC

Fig.1(b) to (e) show the 3D scatter plot of 291 charged groups in the region around the PTC and the tunnel entry port. The exact number of phosphate moieties around the PTC was counted from the x-ray solved structural data. The *bare* surface charge density *σ*\* was calculated by dividing the number of phosphate moieties (= 100) closer than 12 *Å* from the truncated spheroid by its surface area. The surface area was analytically calculated to be 15.75 nm^2^ as detailed in the appendix 6. Hence, the *bare* surface charge density *σ*\* is estimated to be 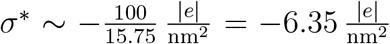 on the surface of the spheroid cavity around the PTC. This numerical result for the surface charge density is approximately three times higher than the surface charge density prevailing on the inner surface of the ribosome exit tunnel [15].

It is unclear whether Bjerrum or Debye theory should be used for the electrostatic screening length in the medium inside the PTC cavity. The Bjerrum screening length as computed from Eq.(20) is 0.72 nm when water permittivity *∈* = 78 is used at 298 K inside the truncated spheroid medium around the PTC. The Debye screening length as computed from Eq.(21) is 1.1 nm when water permittivity *∈* = 78 and an ionic strength *I* = 75 mM*/*L (= 75 M*/*m^3^) are used at 298 K inside the truncated spheroid medium around the PTC. Note that a Debye screening length equal to the Bjerrum length corresponds to the ionic strength’s value of *I* = 76 mM*/*L at 25°C and for water permittivity. The mathematical expressions of both theories are then numerically equivalent and knowing which theory prevails is numerically irrelevant.

The relative permittivity prevailing in the medium around the PTC is not known with accuracy and is not homogeneous. The medium inside the PTC cavity is more aqueous than the medium in the more confined microenvironment of the ribosome exit tunnel. It was hypothesized in this study that a coarse grained permittivity should be taken in the range corresponding to a mixture of protein and water, i.e at least between *∈*_*r*_ = 8 and *∈*_*r*_ = 78.

### 4.2. Comparison of the potential profiles and fields calculated from the spheroid idealized shape and from the x-ray solved structural data around the PTC

The x-ray solved structure dataset includes the exact 3D coordinate positions of a total of 291 charged atoms that are closest to the ribosome exit tunnel entry point or the PTC. Of these 291 atoms, 113 are at a distance shorter than 12 *Å* from the truncated spheroid’s outer surface. The volume inside the truncated spheroid is empty, i.e., there are no 23SrRNA phosphorus atoms or ribosomal proteins charged amino acid moieties inside the volume. The charges which are at a distance larger than 12 *Å* from the outer surface of the cavity are so strongly screened by the dielectric medium that they can be neglected as was detailed in [15].

The electrostatic potential profile Φ(*z*) for the uniformly charged idealized spheroidal shape, as numerically calculated from Eq.(16), is shown in Fig.2(c) black solid line. Upon implementing Eq.(19) in Python and using the exact positions of the 113 charged atoms within 12 *Å* of the spheroid’s outer surface as mapped in Fig.1, we obtained the electrostatic potential along the spheroid’s centerline shown in Fig.2. The electrostatic potential contributed by the phosphate moieties, Fig.2(a) is negative while the potential contributed by the charged amino acid residues Fig.2(b) is positive as arginine or lysine outnumber aspartate or glutamate residues. The net resulting electrostatic potential is negative, orange solid line in Fig.2(c,) as there are ∼ 100 phosphate groups and only ∼ 13 charged amino acid residues at a distance closer than 12 *Å* from the PTC surface cavity. The charged amino acid residues are located much further away from the catalytic center than are the phosphate moieties. The potential profile is due to the dominant presence of negatively charged phosphate moieties harboured by 23S/28S rRNA on the inner surface cavity in the immediate vicinity of the PTC.

The potential profiles calculated from the uniformly charged spheroid idealized shape (Eq.16) or calculated from the discrete sum of the Yukawa-Debye-Huckel formula over the charged atoms at their exact positions (Eq.19) are compared in Fig.2(c). The two potential profiles look similar and provide an estimation of the order of magnitude of the electrostatic potential along the virtual centerline between the PTC and the entry port of the exit tunnel. The uniformly charged assumption of the surface in the region between the PTC and the tunnel entry port appears to be reasonable. The discrepancy between the two potential profiles (idealized spheroidal shape with uniform surface charge density versus exact atomic positions of the charges), Fig.2(c), or axial electrostatic force profiles, Fig.2(d), is sensitive to the two phenomenological parameters of the models i.e., permittivity and screening length and their possible local heterogeneity (not shown here). Actual measurements of the electric field through vibrational Stark effect spectroscopy would provide indirect support and constraints on the ranges of the local values for *∈* and *ξ* (permittivity and screening length). The electrostatic potential particular profile results from the funnel shape of the cavity.

The important feature of the funnel shape on the potential profile is that the potential is smoothly convex (smaller curvature) and then sharply convex (larger curvature) when moving from the PTC to the tunnel entry port along the *z*−axis, Fig.2(c). The resulting negative inverse bell shape peak for the electric field (or force) has a larger width in the region from the PTC to the tunnel entry port than the width of the positive bell shape peak in the region near the tunnel entry port, Fig.2(d). The respective electric fields profiles and hence the forces along the *z*−axis centerline are also compared in Fig.2(d). From these numerically estimated electrostatic force profiles, the maximum force at the center of the cavity neighbouring the PTC would be between −21 and −27 pN for a unit positive test charge. The negative sign means that the force would point from the PTC to the tunnel entry port for a positive test charge.

### 4.3. Electric field estimation in the vicinity of the PTC

The ratio of the *bare* surface charge density over the medium permittivity *σ*\**/∈* has the dimension of an electric field. In the absence of a screening layer of water molecules constitutive of the cavity inner wall, the order of magnitude of this ratio in the immediate vicinity of the PTC cavity surface is 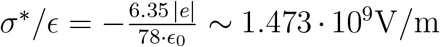, i.e., = 14.73 MV*/*cm, at least if the cavity medium is fully filled with water. If the medium in the PTC is only partially filled with water (*∈* = 78), and the main components are ribonucleic acids (tRNAs) (*∈* = 8) and the carboxy terminal end of the growing nascent protein (*∈* = 4); then the resulting medium coarse grained permittivity would be *∈* ∼ 8 (e.g. for a medium composed of 5% water and 95% protein). In this latter case, the estimated electric field, or the ratio 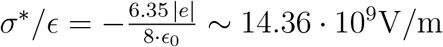, i.e., = 143.6 MV*/*cm.

Using vibrational Stark effect spectroscopy, Fried, Boxer and coworkers measured the electric field in a typical enzyme-substrate configuration at the catalytic site of the proteic enzyme ketosteroid isomerase (KSI), with a magnitude of 144 ± 6 MV*/*cm [5, 21]. Our numerical result shows that the order of magnitude of the electric field in the vicinity of the catalytic surface of the peptidyl transferase center of the ribosome is similar to that of the catalytic sites of known protein enzymes.

This intense electric field is possibly and locally electrostatically screened by constitutive water molecules of the PTC cavity inner wall. The reorientation of the permanent electric dipoles of constitutive water molecules on the inner surface of the PTC cavity would strongly damp the electrostatic potential and give rise to an apparent surface charge density 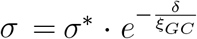 at a distance *δ* from the support of the *bare* charges, where *ξ*_*GC*_ = 0.105 nm is the Gouy-Chapman length. The Gouy-Chapman length is the screening length used when dealing with charges distributed on a surface wall and when the screening is mainly due to water molecules (not mobile ions) [15, 20].

The apparent electric field along the *z*-axis as numerically calculated from the negative of the first derivative of Eq.(16), Fig.2(d) black solid line, or of Eq.(19), Fig.2(d) black dashed line, is approximately equal to 1.3 MV*/*cm. The corresponding axial force experienced by a positively charged unit test probe would be 21 − 27 pN around the PTC region as shown on Fig.2(d).

The shape of the prolate spheroid with a uniformly charged inner surface results in a particular profile for the electrostatic potential and for the resultant electric field along the centerline of the cavity towards the tunnel entry point.

## 5. CONCLUSIONS

End-users of structures are functional biologists and enzymologists. The salient feature of the catalytic center of the ribosomal large subunit is that it is made only of nucleic acids. This is uncommon as the catalytic activity in biochemical processes are carried out mostly by proteins. In this study, it was shown that structural data can be used to provide insights on the electrostatic environment around the catalytic center of the ribosome where the peptide bond is formed and in the cavity towards the tunnel entry point where the nascent protein is threaded.

In future perspective, we expect that the quantitative estimation of the electrostatic potential and field around the PTC will be useful to shed light on the kinetics of the peptide bond formation and on its dependence of the nature of the newly incorporated amino acid residues. More precisely, the quantitative knowledge of the electric field will allow to calculate the mechanical work exerted by the field along any displacement of the newly incorporated amino acid in the nascent growing peptide. The activation energy of the peptide bond formation will be increased or decreased depending on the sign of this mechanical work.

### 6 APPENDIX

#### 6.1. Analytical solution for the area of the truncated prolate spheroid as a surface of revolution of a truncated ellipse

A prolate spheroid is a surface *S* ∈ ℝ^3^ generated by the revolution of an ellipse about its major axis. A parametric representation of a simple ellipse in the plane *Oxy* in ℝ^3^ having semi-major axis *a* and semi-minor axis *b* is

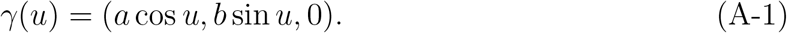

The parametric representation of the surface of revolution is

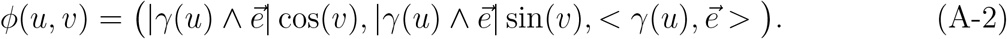

for all (*u, v*) ∈ *K*, with *K* = [*u*_*lower*_, *u*_*upper*_] × [0, 2*π*] and 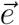 the unit vector about which the revolution takes place. The area of the surface is determined by the formula known in elementary mathematical analysis

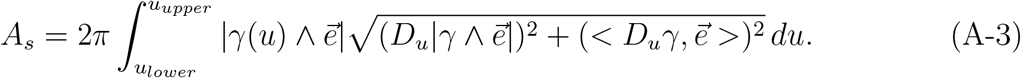

Taking the revolution about the semi-major axis, we have 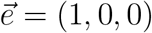 and

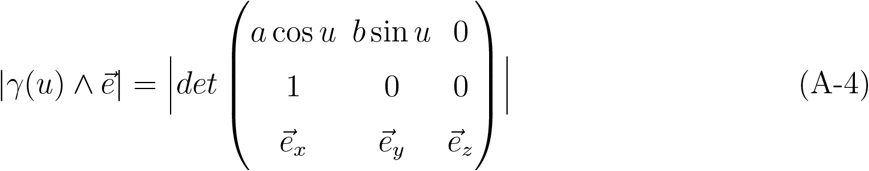

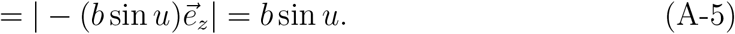

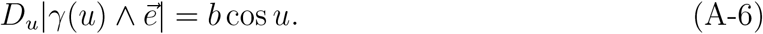

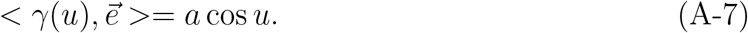

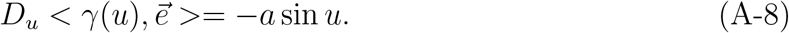

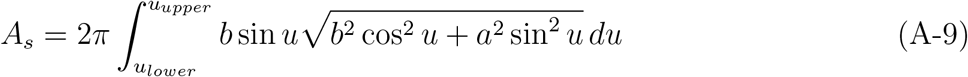

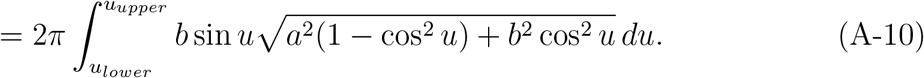

Substituting *t = cos u, dt = −sin u du*, we have

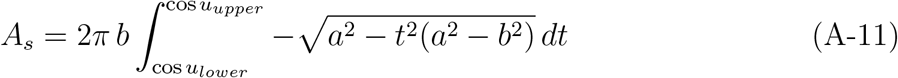

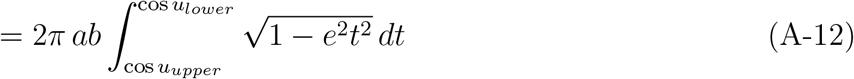

where, in the previous line, the eccentricity of the ellipse 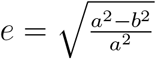 was used. Substituting again *et* = sin *w, e dt* = cos *w dw*, the last integral turns into

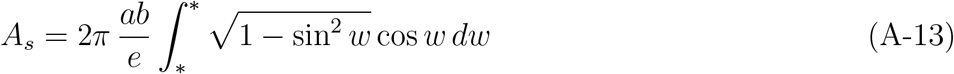

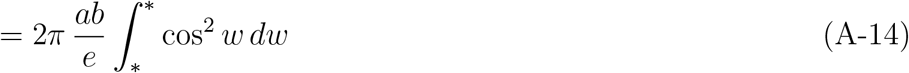

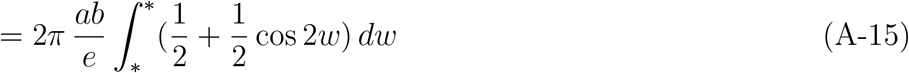

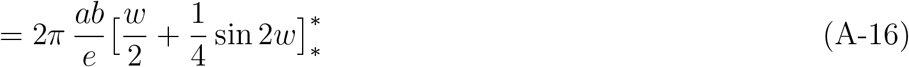

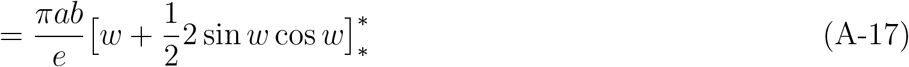

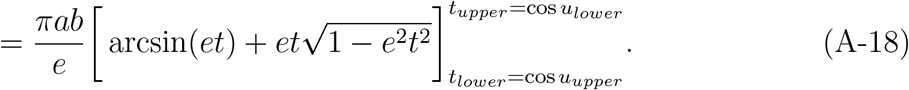

The area of the truncated prolate spheroid of semi-major axis *a* = 8.9744 nm and semi-minor axis *b* = 2.25 nm (eccentricity *e* = 0.9681), for *t* in the range *t*_*lower*_ = cos *u*_*upper*_ = 0.7454 and *t*_*upper*_ = cos *u*_*lower*_ = 0.9750, is calculated from this last formula to be 15.75 nm^2^. Note that, when 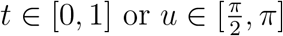, the surface of the half prolate spheroid is 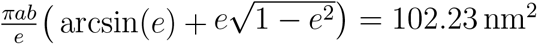 with the adopted values of *a* and *b*.

## ACKNOWLEDGMENTS

This work was supported by the FNRS-FWO EOS grant *n*^*o*^30480119 (Join-t-against-Osteoarthritis), the FNRS-WELBIO grant *n*^*o*^*WELBIO* − *CR* − 2017*S* − 02 (THERAtRAME) in Belgium and the European Research Council under the European Union’s Horizon 2020 Framework Program (H2020/2014-2020) /ERC grant agreement *n*^*o*^772418 (INSITE).

